# Bacteria-mediated stabilization of a panel of Picornaviruses

**DOI:** 10.1101/410167

**Authors:** Elizabeth R. Aguilera, Y Nguyen, Jun Sasaki, Julie K. Pfeiffer

## Abstract

Several viruses encounter various bacterial species within the host and in the environment. Despite these close encounters, the effects of bacteria on picornaviruses specifically is not completely understood. Previous work determined that poliovirus (PV), an enteric virus, has enhanced virion stability when exposed to bacteria or bacterial surface polysaccharides such as lipopolysaccharide. Virion stabilization by bacteria may be important for inter-host transmission since a mutant PV with reduced bacterial binding had a fecal-oral transmission defect in mice. Therefore, we investigated whether bacteria broadly enhance stability of picornaviruses from three different genera: *Enterovirus* (PV and coxsackievirus B3 (CVB3)), *Kobuvirus* (Aichi virus) and *Cardiovirus* (Mengo virus). Furthermore, to delineate strain-specific effects, we examined two strains of CVB3 and a PV mutant with enhanced thermal stability. We determined that specific bacterial strains enhance thermal stability of PV and CVB3, while Mengo virus and Aichi virus are stable at high temperatures in the absence of bacteria. Additionally, we determined that bacteria or lipopolysaccharide can stabilize PV, CVB3, Aichi virus, and Mengo virus during exposure to bleach. These effects are likely mediated through direct interactions with bacteria since viruses bound to bacteria in a pull-down assay. Overall, this work reveals shared and distinct effects of bacteria on a panel of picornaviruses.

**IMPORTANCE:** Recent studies have shown that bacteria promote infection and stabilization of poliovirus particles, but the breadth of these effects on other members of the *Picornaviridae* family is unknown. Here, we compared the effect of bacteria on four distinct members of the *Picornaviridae* family. We found that bacteria reduced inactivation of all of the viruses during bleach treatment, but not all viral strains were stabilized by bacteria during heat treatment. Overall, our data provide insight into how bacteria play differential roles on picornavirus stability.

## INTRODUCTION

The *Picornaviridae* family includes important human pathogens that can cause a range of diseases such as the common cold, meningitis, hepatitis, and paralysis. The *Picornaviridae* family is diverse and currently includes 80 species in 35 genera. Members of this family are nonenveloped and contain a single-stranded, positive-sense viral genome approximately 7,500 nucleotides in length (1).

Recent studies have shown that bacteria, in particular the gut microbiota, play several important roles during viral infection. Enteric viruses encounter a milieu of microorganisms, including bacteria, both within and outside of the host. It is estimated that these viruses encounter approximately 10^11^ bacteria in the host and are expected to encounter even more in the environment (2). Indeed, bacteria enhance infection of several unrelated viruses, including poliovirus, reovirus, rotavirus, mouse mammary tumor virus, and norovirus (3-9). These “pro-viral” effects are mediated by two known mechanisms: 1) Direct interactions between bacteria and viruses that increased virion stability and attachment to host cells, and 2) indirect interactions between bacteria and the host immune system that modulated immune responses for productive viral infection (3-8,10).

Intriguingly, bacteria and bacterial molecules can inhibit infection with certain viruses. For example, Ichinohe *et al*. demonstrated that certain bacteria promote host immune responses during influenza infection of mice (11). More recently, Bandoro *et al*. determined that exposure to the bacterial surface molecule lipopolysaccharide (LPS) reduced stability of several strains of influenza virus by altering the morphology of the virion envelope (12).

Based on the importance of bacterial-viral interactions on viral infection, we sought to determine whether bacteria differentially affect different members of the same viral family, the *Picornaviridae*. We used a panel of four picornaviruses that are spread by the fecal-oral route and represent three separate genera: *Enterovirus* (coxsackievirus B3 (CVB3) and PV), *Kobuvirus* (Aichi virus) and *Cardiovirus* (Mengo virus). We found that a subset of the viral panel were stabilized by bacteria during heat treatment but that all of the picornaviruses tested were stabilized by bacteria during bleach treatment. We also determined that viruses bound to bacteria, indicating that direct interactions may be facilitating viral stabilization of these viruses. This work expands on bacteria-mediated enhancement effects previously observed with PV to other members of the *Picornaviridae* family. Ultimately, this work defines the unique interactions between specific viruses and bacteria which may provide insight into virion environmental stability and transmission.

## RESULTS

### Panel of viruses from the *Picornaviridae* family and bacterial strains

Previous studies have indicated that bacteria can reduce the inactivation of poliovirus particles after heat or bleach treatment (3,7). In order to investigate whether bacteria stabilize other members of the *Picornaviridae* family from these inactivating conditions, we selected viruses from separate genera and viruses with differences in capsid sequence similarity (**Fig. 1 A, B and Table 1**). These viruses differ in their capsid structure sequence and topology, which may confer different interactions with bacteria (**Fig. 1A**). The panel is composed of one virus from the *Kobuvirus* genus (Aichi virus), one virus from the *Cardiovirus* genus (Mengo virus) and three viruses from the Enterovirus genus (PV, CVB3-H3 and CVB3-Nancy). CVB3-Nancy and CVB3-H3 have 98.4% capsid sequence similarity at the amino acid level and were compared to determine whether there are strain-specific differences in bacteria-mediated stabilization (**Fig. 1B and Table 1**). Additionally, our panel included a PV mutant with a single amino acid change in the VP1 capsid coding region (PV-M132V), that confers enhanced thermal stability in the absence of bacteria (13).

**Figure 1.**
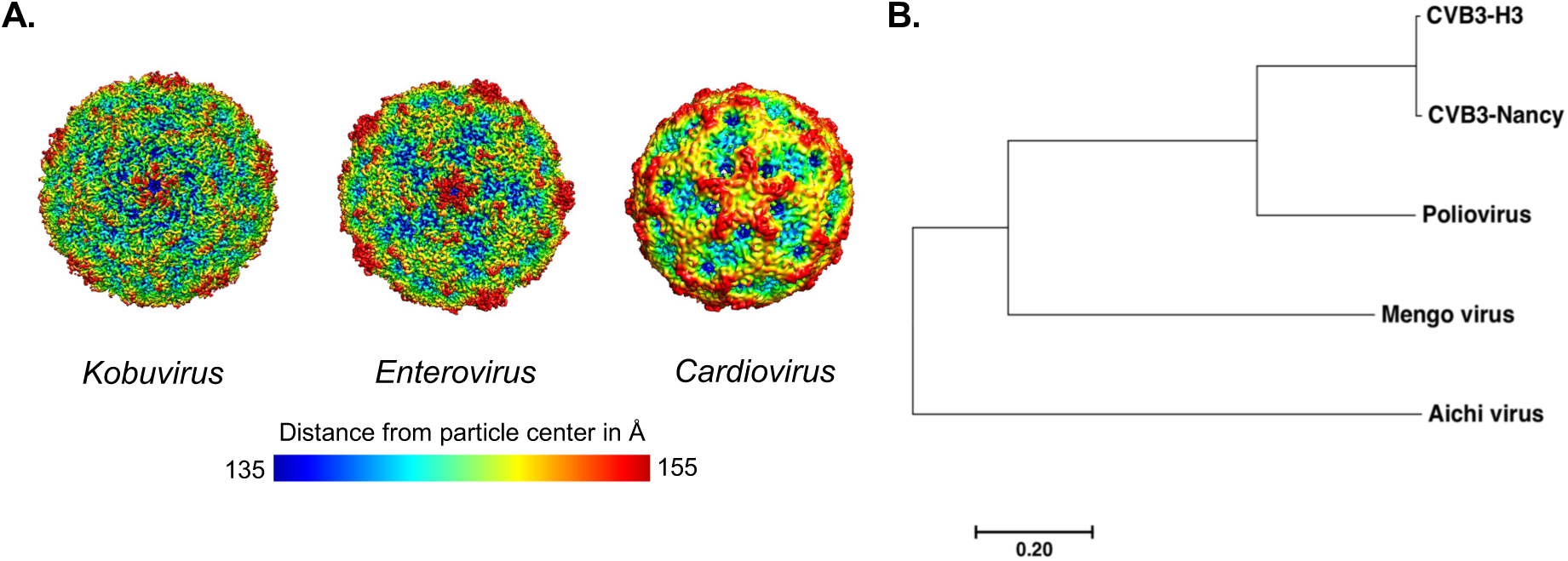
Panel of picornaviruses used in this study. **A**) Structural models of picornaviruses. Structural comparisons were performed using EMDB ascension numbers for each viral genus and topological distances from the center of the virion calculated from 135 Å (blue) to 155 Å (red), as indicated by the scale bar (22). Representative viruses for each genus are Aichi virus (*Kobuvirus*), CVB3 (*Enterovirus*), and Saffold virus (*Cardiovirus*). The Aichi virus structure is at 3.7 Å resolution, CVB3 structure is at 3.9 Å resolution and Saffold virus is at 10.6 Å resolution. Models and distances were generated with UCSF Chimera software. **B**) Phylogenetic tree of picornaviruses based on the amino acid sequence of the capsid-coding region. The tree was generated using MEGA7 software. The evolutionary history was inferred using the Neighbor-Joining method. The scale bar represents the number of substitutions per site.

We also selected a representative panel of enteric bacteria, and bacterial and non-bacterial molecules (**Table 2**). We included LPS, which is glycan found on the surface of Gram negative bacteria. Additionally, we examined two representative Gram negative bacterial strains (*Escherichia coli* 1470 and *Prevotella ruminicola)*, and two representative Gram positive bacterial strains (*Bacillus badius* and *Lactobacillus johnsonii).* We previously showed that *E. coli* 1470, *P. ruminicola, B. badius* and *L. johnsonii* bind to PV (14), but whether these strains stabilize PV and other picornaviruses was unknown. We also previously showed that non-bacterial compounds, such as bovine albumin serum (BSA) and cellulose, had minimal effects on PV stability and were included in this study as controls (**Table 2**) (3,7).

**Table 1.** Percent sequence identity between panel of picornaviruses.

| Capsid Amino Acid Sequence Similarity (%) |  |  |  |  |  |  |
| --- | --- | --- | --- | --- | --- | --- |
| Family and genus | Virus | Aichi virus | Mengo virus | CVB3 H3 | CVB3 Nancy | Poliovirus |
| *Picornaviridae* |  |  |  |  |  |  |
| *Kobuvirus* | Aichi virus | 100.0 | 28.6 | 23.3 | 23.3 | 24.3 |
| *Cardiovirus* | Mengo virus |  | 100.0 | 30.0 | 29.8 | 29.0 |
| *Enterovirus* | CVB3 H3 |  |  | 100.0 | 98.6 | 54.4 |
|  | CVB3 Nancy |  |  |  | 100.0 | 54.2 |
|  | Poliovirus |  |  |  |  | 100.0 |

**Table 2.**
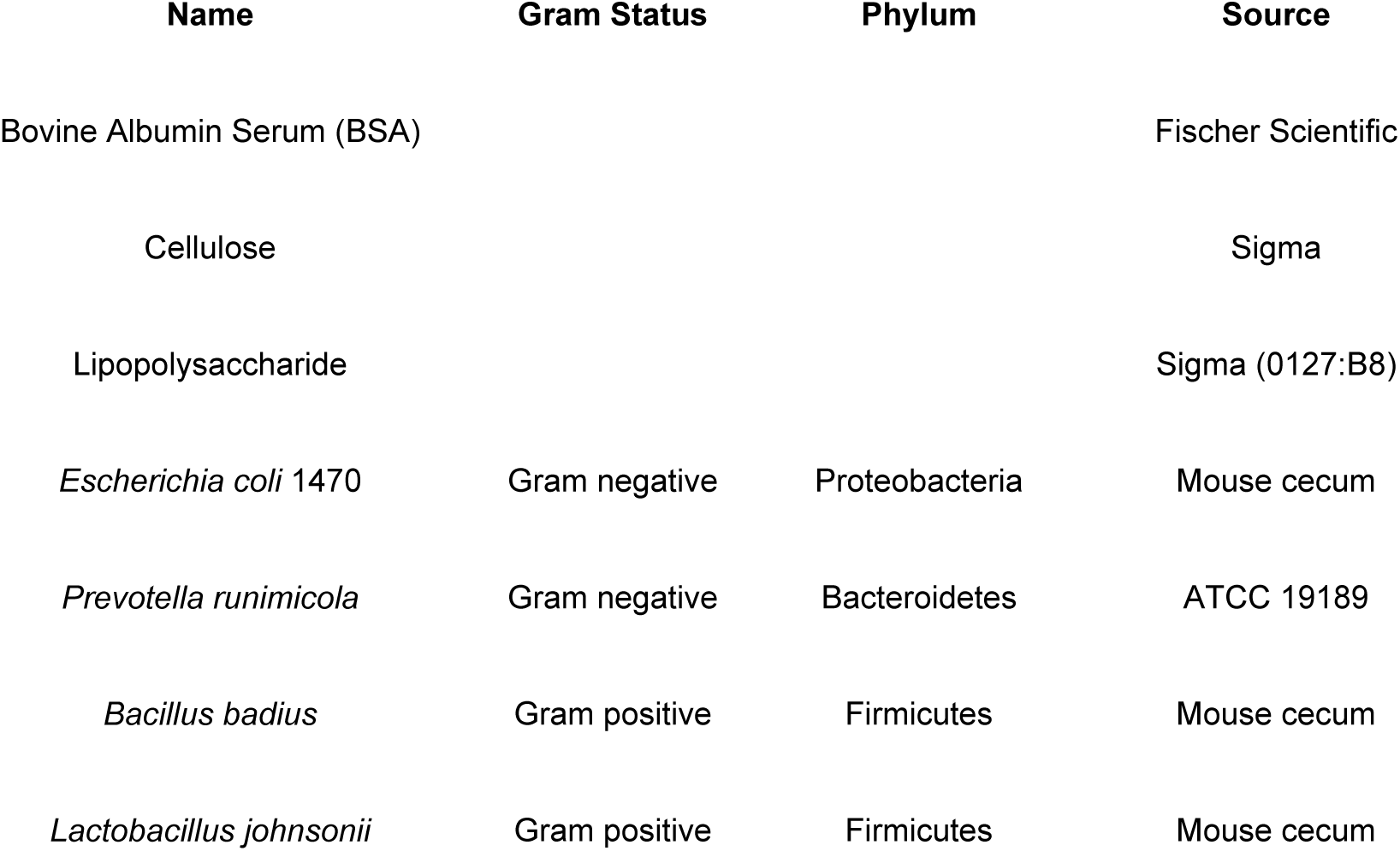
List of reagents and bacterial strains used in this study.

| Name | Gram Status | Phylum | Source |
| --- | --- | --- | --- |
| Bovine Albumin Serum (BSA) |  |  | Fischer Scientific |
| Cellulose |  |  | Sigma |
| Lipopolysaccharide |  |  | Sigma (0127:B8) |
| *Escherichia coli* 1470 | Gram negative | Proteobacteria | Mouse cecum |
| *Prevotella runimicola* | Gram negative | Bacteroidetes | ATCC 19189 |
| *Bacillus badius* | Gram positive | Firmicutes | Mouse cecum |
| *Lactobacillus johnsonii* | Gram positive | Firmicutes | Mouse cecum |

### Specific bacteria enhance stability of a subset of picornaviruses during heat treatment

To determine whether bacteria enhance stability of picornaviruses, we first examined viral inactivation at elevated temperatures. Picornavirus particles can be inactivated by undergoing premature genome release at a range of temperatures, with faster inactivation at higher temperatures (7,15,16). To increase tractability of our assays, we used relatively high temperatures for our thermal inactivation experiments because inactivation occurs relatively quickly. We first tested viral stability at 44°C for 4.5 h, a condition that we determined inactivates approximately 99% of PV infectivity during incubation in PBS (**Fig. 2A**). Viruses were mixed with PBS, compounds (BSA, Cellulose or LPS), or bacteria, incubated at 44°C for 4.5 h, and plaque assays were performed to quantify the amount of viable virus remaining. When we incubated PV with any of the bacterial strains or LPS, we observed >50-fold increase in viral stability compared to PBS (**Fig. 2A**). A similar stabilization was observed for CVB3-H3 and CVB3-Nancy when compared to PBS. Non-bacterial compounds (BSA and Cellulose) did not stabilize PV or either CVB3 strain. Interestingly, Aichi virus and Mengo virus were very stable in PBS under these conditions. We also tested a recently identified heat-resistant PV mutant, PV-M132V (13). Like Aichi virus and Mengo virus, PV-M132V was resistant to heat treatment, and thus incubation with any of the compounds or bacterial strains did not increase stability (**Fig. 2A**).

We next wanted to determine whether bacteria could increase stability of the heat-stable viruses at temperatures where they become heat labile. First, we increased the temperature in the thermal stability assay to 46°C, and found that similar to the 44°C assay, PV, CVB3-H3, and CVB3-Nancy were stabilized by bacteria (**Fig. 2B**). Aichi virus, Mengo virus, and PV-M132V were still stable in the 46°C assay and incubation with any of the bacteria compounds or strains did not increase stability. To determine the temperature necessary to inactivate Aichi virus and Mengo virus, we tested viability at temperatures from 46-58°C for 4.5 h. These additional experiments at different temperatures revealed that Aichi virus and Mengo virus were ~99% inactivated when incubated at 50°C or 57°C for 4.5 h, respectively (**Fig. 2C-2F**). Despite viral inactivation during these conditions, none of the bacterial strains or bacterial polysaccharides could stabilize either of these viruses (**Fig. 2D and 2F**). Intriguingly, BSA stabilized Aichi virus during incubation at 50°C, indicating this virus may have different requirements for stabilization during heat treatment (**Fig. 2D**). Overall, these data indicate that bacteria do not stabilize Aichi and Mengo virus during incubation at high temperatures, but that bacterial stabilization of these viruses may be less important given their inherent high stability.

**Figure 2.**
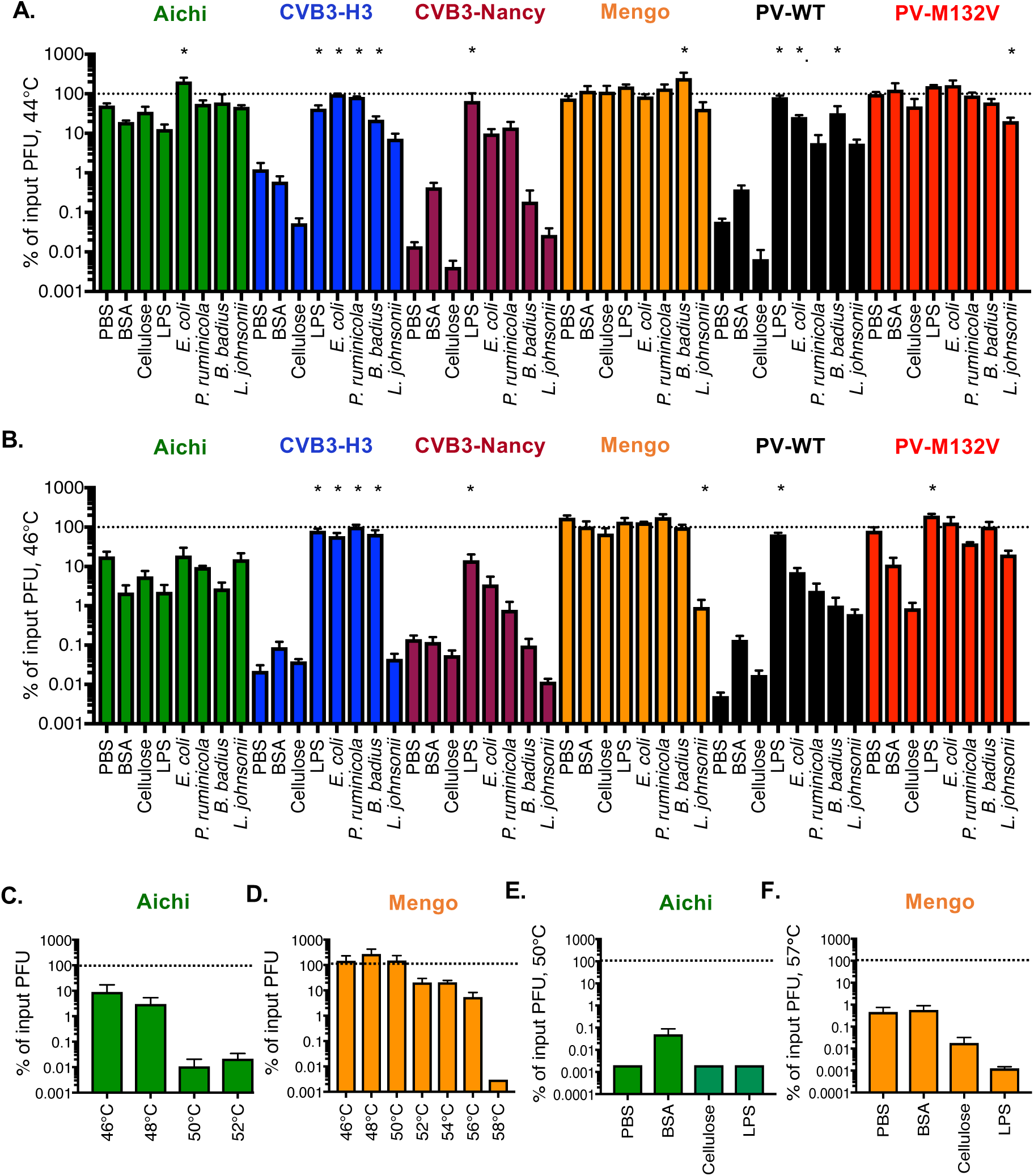
Effects of bacteria and compounds on picornavirus stability at elevated temperatures. Thermal stability assays were performed by incubating 1 × 10^5^ PFU viruses in PBS, 1 mg/mL BSA, cellulose, LPS, or 1 × 10^10^ CFU of bacterial strains at various temperatures for 4.5 h. The amount of viable virus following each assay was determined by plaque assay and compared to PBS viral titer at 0 h to determine percent of input PFU. **A**) 44°C assay. Data are representative of ten to eighteen independent experiments, n= 4-47. **B**) 46°C assay. Data are representative of nine to fourteen independent experiments, n= 4-25. **C**) Incubation of Aichi virus in PBS at various temperatures. Data are representative of two to three independent experiments, n=3-5.**D**) Incubation of Mengo virus in PBS at various temperatures. Data are representative of one to three independent experiments, n=2-6. **E**) Aichi virus 50°C assay. Data are representative of two independent experiments, n=4. **F**) Mengo virus 57°C assay. Data are representative of two experiments, n=4. Bars are shown in SEM. Statistical significance was determined by one-way ANOVA, * = *P*<0.05. n.s., not significant.

### The effect of feces on picornaviruses

We next wanted to determine whether the viruses in our panel are stabilized in feces. As enteric viruses are transmitted by the fecal-oral route, the potential effects of fecal components on their stability and infection is highly relevant. We previously showed that PV is stabilized in feces from conventional mice (7). Here, we compared viral stability when viruses were incubated in PBS or feces from conventional mice over the course of several days at 37°C followed by quantification of remaining viable virus by plaque assay. We found that Aichi virus was not stabilized by feces compared to PBS at 37°C after Day 1 (**Fig. 3A**). Feces only moderately stabilized CVB3-H3 at early timepoints (**Fig. 3B**). However, CVB3-Nancy and PV were stabilized by feces at later timepoints (see Day 8) (**Fig. 3C and 3E**). We also demonstrated that PV-M132V was stable during 8 days of incubation at 37°C in both PBS and feces (13). Interestingly, Mengo virus exhibited significant inactivation after 4 days at 37°C, but incubation in feces limited this inactivation (**Fig. 3D**). This result was surprising given the stability of Mengo virus at 46°C for 4.5 h and the lack of bacterial stabilization of Mengo virus at 57°C (**Fig. 2B and 2F**). However, Mengo virus may have enhanced thermal sensitivity over longer time courses and bacterial effects may be apparent only under these conditions and/or non-bacterial components of feces could affect Mengo virus. When we incubated Mengo virus with mixtures of *E. coli, P. ruminicola*, and *L. johnsonii* at 37°C for several days, Mengo virus was stabilized compared to PBS at 37°C (see dashed line compared to dotted line) (**Fig. 3F**). These findings indicate that bacteria stabilize Mengo virus during longer exposures to body temperature (37°C) (**Fig. 3F**). Overall, these data indicate that several picornaviruses, but not all, are stabilized in feces, which could facilitate transmission.

**Figure 3.**
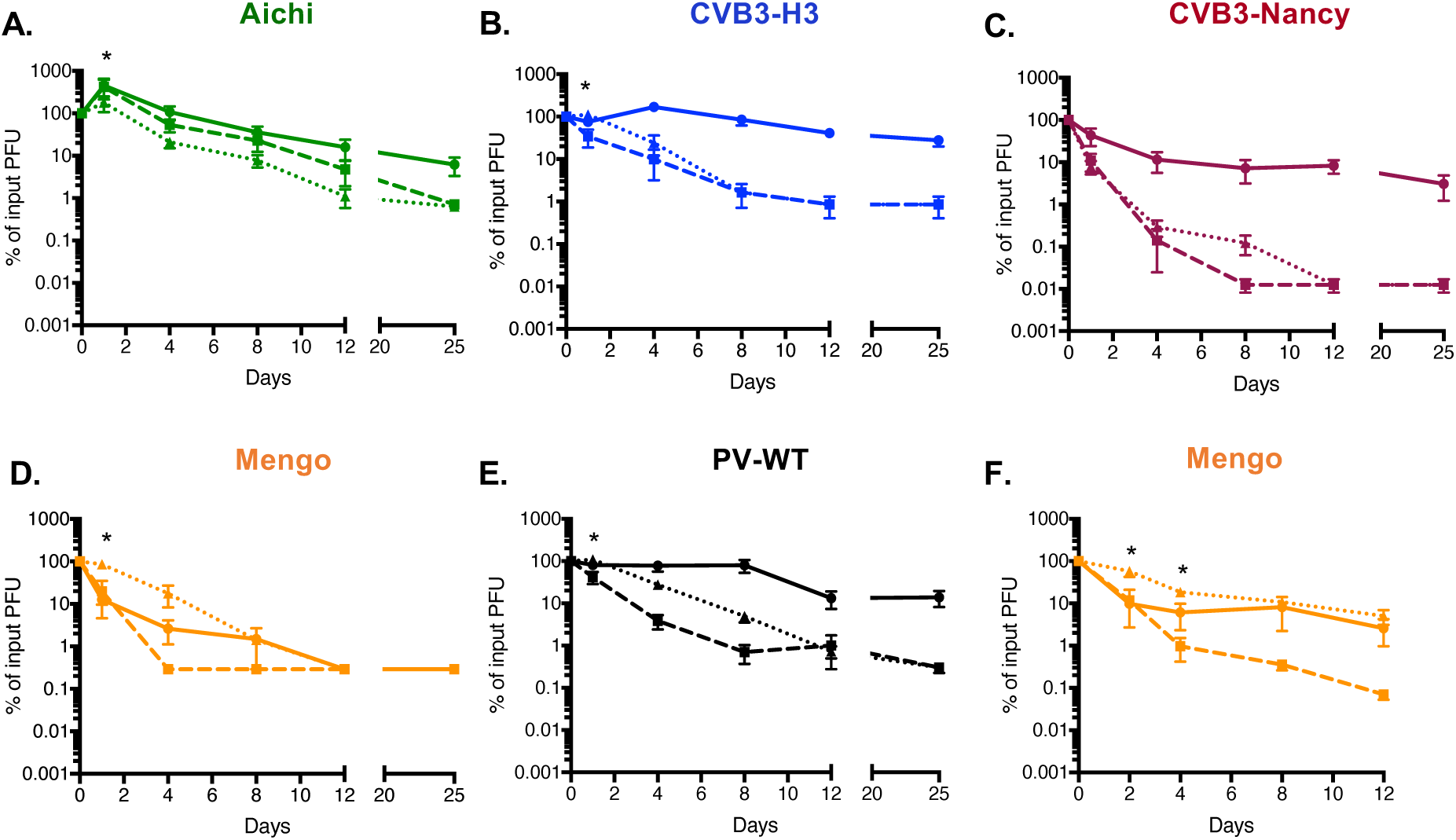
Picornavirus stability in feces. 1 × 10^5^ PFU of A) Aichi virus, B) CVB3-H3, C) CVB3-Nancy, D) Mengo virus, or E) PV was incubated with PBS or a slurry of feces from mice and incubated at 37°C (PBS, dashed lines; Feces, dotted lines) or 4°C (PBS, solid lines). Data are representative of two to three experiments, n = 4-5. In F) 1 × 10^5^ PFU of Mengo virus was incubated with PBS or a mixture of *E. coli, P. ruminicola*, and *L. johnsonii* and incubated at 37°C (PBS, dashed lines; Bacteria, dotted lines) or 4°C (PBS, solid lines). Data are representative of two independent experiments, n = 4. Samples were taken at designated time points and processed prior to plaque assay for quantification of viable virus. Bars are shown in SEM. Statistical significance between PBS and feces or bacteria at 37°C was determined by two-way ANOVA, * = *P*<0.05.

### Bacteria enhance stability of picornaviruses during bleach exposure

In addition to heat, virions can be inactivated by chlorine bleach via capsid penetration and damage and/or genome release (17-21). To determine whether bacteria affect bleach inactivation of viruses, we pre-incubated viruses in PBS, compounds, or bacterial strains for 1 h followed by exposure to dilute bleach (0.0001%) for 1 min, neutralization, and plaque assay to determine the amount of viable virus present. We determined that when pre-incubated in PBS, all viruses lost ~90% of their infectivity (**Fig. 4**). However, when pre-incubated with LPS or bacterial strains, all viruses were stabilized by at least some of the treatments (**Fig. 4**). Importantly, pre-incubation of the viruses with BSA or cellulose did not prevent viral inactivation by bleach treatment, indicating that the effects were specific to bacteria and LPS and not just due the presence of additional molecules (**Fig. 4**). Interestingly, the heat stable PV-M132V mutant virus was inactivated by bleach to the same extent at PV-WT, and bacteria limited bleach inactivation of PV-M132V. These results suggest that thermal inactivation and bleach inactivation occur through separable mechanisms, and that bacteria stabilize virions for both. Overall, these results indicate that bacteria enhance viral stability of fecal-orally transmitted picornaviruses during bleach treatment.

**Figure 4.**
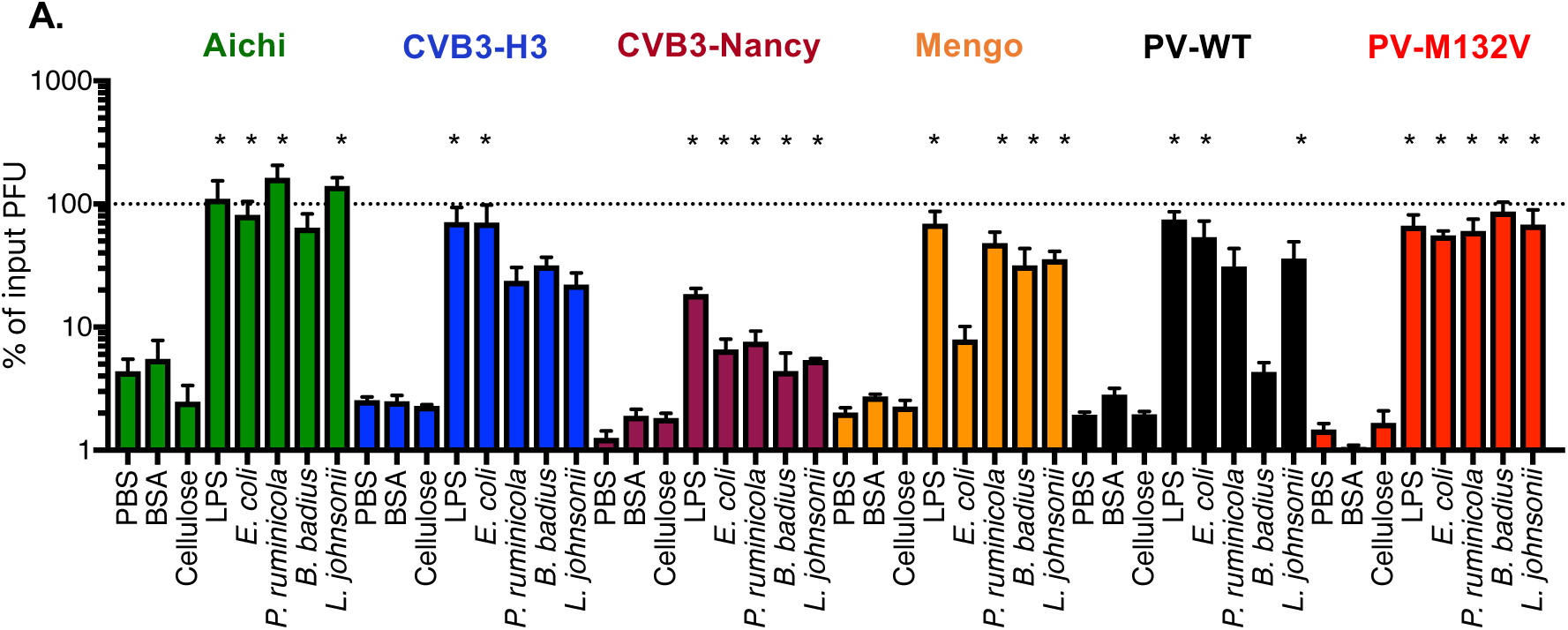
Effects of bacteria on picornavirus stability during bleach treatment. 1 × 10^5^ PFU viruses were incubated individually in PBS, 1 mg/mL BSA, cellulose, LPS, or 1 × 10^8^ CFU of bacterial strains at 37°C for 1 h. After incubation, samples were treated with 0.0001% bleach for 1 minute and neutralized with sodium thiolsulfate. Amount of viable virus was determined by plaque assay and compared to PBS viral titer at 0 h to determine % of input PFU. Data are representative of seven to twenty independent experiments, n = 4-40. Bars are shown in SEM. Statistical significance was determined by one-way ANOVA, * = *P*<0.05. n.s., not significant.

### Bacteria bind to a select panel of picornaviruses

Since bacteria enhanced stability of specific picornaviruses during heat or bleach inactivation, we wanted to determine whether viruses directly interact with bacteria. In particular, we were curious whether bacterial binding efficiencies vary among closely related viruses, such as CVB3-Nancy and CVB3-H3, or between the PV-M132V heat stable mutant and PV-WT. Previously, we showed that PV can bind directly to the surface of bacteria (3,7,14). ^35^S-labeled CVB3-Nancy, CVB3-H3, PV-WT, or PV-M132V were incubated with beads, *B. badius*, or *E. coli* for 1 h followed by centrifugation, washing, and scintillation counting the bacterial pellets to quantify viral binding. We determined that PV-WT and PV-M132V bound to both bacterial strains to approximately the same extent (**Fig. 5A**). This indicates that while the PV-M132V mutant does not require the presence of bacteria for stability during heat treatment, it still binds to bacteria, which could explain why bacteria limit bleach inactivation of PV-M132V (**Fig. 4**). Additionally, we determined that both CVB3 strains bind to the two bacterial strains tested (**Fig. 5B**). Interestingly, binding of CVB3-Nancy to *E. coli* was nearly 3-fold higher than CVB3-H3 (**Fig. 5B**). Overall, these results indicate that multiple picornaviruses bind to bacteria, but with different efficiencies.

**Figure 5.**
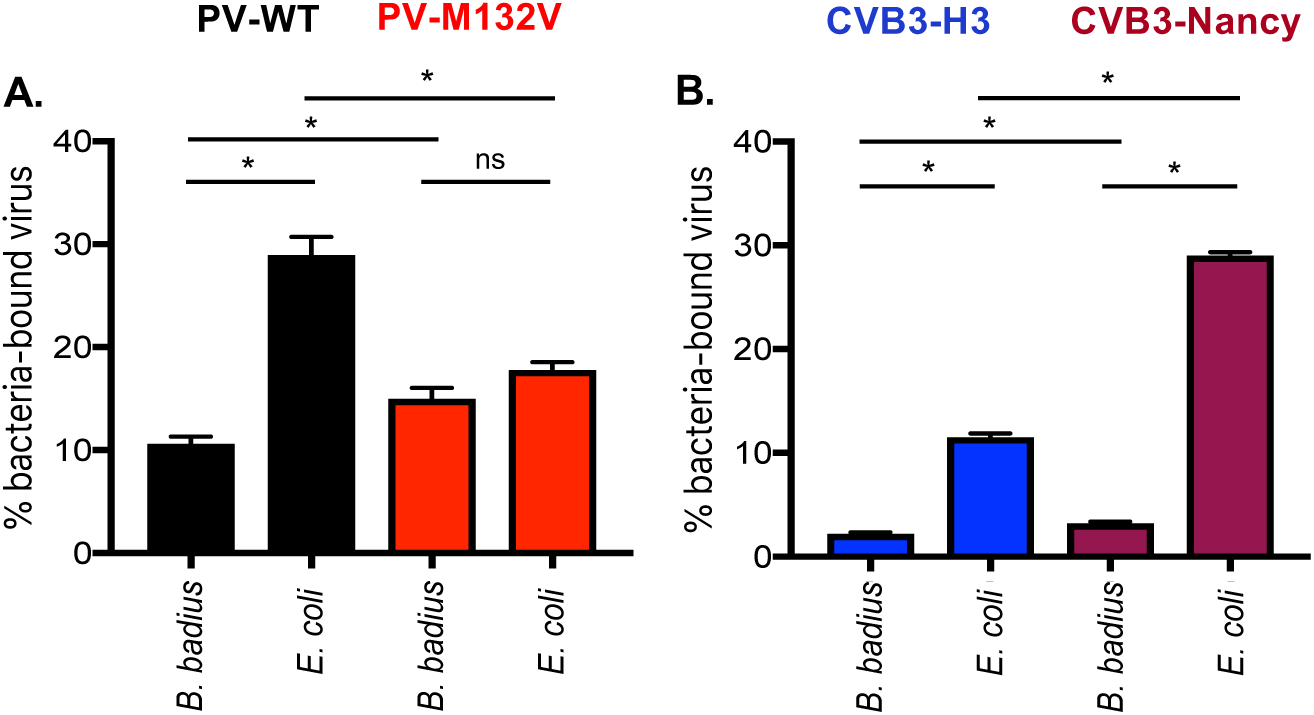
Picornaviruses bind to bacteria. ^35^S-labeled viruses (3,000 CPM/approximately 1 × 10^6^ PFU) were incubated with 1 × 10^8^ CFU of bacteria for 1 h at 37°C. After incubation, samples were spun down and washed to remove unbound virus. Bound virus was quantified by scintillation counting. Data are representative of two independent experiments, n=3-4. Bars are shown in SEM. * = *P*<0.05, based on student’s T test.

## DISCUSSION

The *Picornaviridae* family is diverse and includes a large number of medically relevant human pathogens. While it has been shown that bacteria promote infection, co-infection, and transmission of poliovirus, the impact of bacteria on other picornaviruses is unclear (3,7,14). Here, we show that bacteria increase stability of several viruses from the *Picornaviridae* family, likely through direct interactions.

Our data show that bacteria-mediated thermal stability can vary among a family of viruses. We determined that certain picornaviruses (*Enterovirus* genus members: CVB3-H3, CVB3-Nancy and PV) are sensitive to heat treatment and that bacteria increase stability of these viruses (**Fig. 2 and Fig. 3**). We also determined that another picornavirus (*Cardiovirus* genus: Mengo virus) has mixed phenotypes depending on the condition tested. While Mengo virus was very stable at high temperatures during relatively short incubation times (4.5 h) and was not impacted by bacteria under these conditions, it was inactivated after 4 days at 37°C and exposure to feces reduced this inactivation (**Fig. 3D**). This suggests that Mengo virus may be stabilized by bacteria at physiological temperatures in the host. Finally, we determined that a distantly related picornavirus (*Kobuvirus* genus: Aichi virus) is relatively resistant to high temperature, but is not stabilized by bacteria or bacterial products (**Fig. 2 and Fig. 3A**). In fact, exposure to feces slightly reduced Aichi virus infectivity (**Fig. 3A**). Although Aichi virus is transmitted by the fecal-oral route, there are large sequence and structural differences between Aichi and other picornaviruses that may contribute to the different phenotype (**Fig. 1 A and B**) (22,23). Although a member of the *Caliciviridae*, human norovirus can bind to and is stabilized by bacteria that express certain histo-blood group antigens (24,25). Similarly, reovirus (*Reoviridae* family) can be stabilized by exposure to certain bacteria or bacterial surface molecules, but stabilization efficiency and specificity varies among different reovirus strains (8). Taken together, these results indicate that viruses from separate viral families can be stabilized by bacteria, but that not all viruses within a given family share phenotypes.

While picornaviruses vary in bacteria-mediated thermal stabilization, we found that bacteria enhanced viability of all picornaviruses tested during bleach treatment (**Fig. 4**). Although the PV-M132V mutant was not inactivated at high temperatures, it was inactivated by bleach treatment and bacteria limited this inactivation. Indeed, the PV-M132V virus was determined to bind to bacteria, which could explain stabilization during bleach treatment (**Fig. 4**). Thus, heat inactivation and bleach inactivation are independent and could have separate requirements for stabilization.

Overall, this study provides insight into the effects of bacteria on a panel of viruses from the same family, the *Picornaviridae*. Understanding the role of bacteria during stabilization and infection of viruses could provide insight into efficient infection within specific hosts (i.e. harboring specific microbiota) as well as between hosts (i.e. environmental bacteria).

## MATERIALS AND METHODS

### Cells and viruses

HeLa cells were propagated in Dulbecco’s modified Eagle’s medium (DMEM) supplemented with 10 % calf serum and 1 % penicillin/streptomycin. HeLa cells were used for CVB3, Mengo virus, and PV propagation and quantification of viral titer by plaque assay (26-28). Vero cells were propagated in DMEM supplemented with 10% fetal bovine serum (FBS) and 1 % penicillin/streptomycin. Vero cells were used for Aichi virus propagation and quantification of viral titer by plaque assay. All infections were performed using viruses derived from infectious cDNA clones (the Mengo virus clone was a kind gift from Marco Vignuzzi) (29,30). All viruses were confirmed by Sanger sequencing.

To quantify virus, plaque assay was performed as previously described (26,27,30) Briefly, virus was diluted in phosphate-buffered saline supplemented with 100 μg/mL CaCl_2_ and 100 μg/mL MgCl_2_ (PBS+) and added to cells for 30 min at 37°C in presence of 5 % CO_2_ to allow for attachment. Agar overlay containing DMEM, supplemented with 20 % calf serum, and 2 % agar was used for CVB3 and PV samples and removed after 48 h. Agar overlay containing DMEM, supplemented with 20 % FBS, and 2 % agar was used for Aichi virus samples and removed after 48 h. Agar overlay containing P5 buffer and 2 % agar was used for Mengo virus samples and removed after 48 h (27).

Radiolabeling of picornaviruses was performed as previously described (3,7,28). Briefly, viruses were propagated in the presence of ^35^S-Cysteine/Methionine and were purified using cesium chloride gradients. Purity of viruses were confirmed by phosphorimaging of radiolabeled capsid proteins on SDS-PAGE, and scintillation count to determine CPM and viable fractions (3).

### Bacterial strains

Strains of bacteria were from ATCC or from the cecum of mice, as previously described (14). Cultures were inoculated from glycerol stocks in strain-specific nutrient media as previously described (14). Briefly, cultures were grown overnight, bacterial cell pellets were collected and washed in PBS+. After resuspension in 1 mL PBS+, OD_600_ values were obtained by spectrophotometer (Eppendorf BioPhotometer D30) to determine colony forming units (CFUs) needed specific for each assay. Bacteria were UV inactivated prior to use in assays. The amount of bacteria was confirmed by plating on nutrient-specific agar and conditions prior to UV inactivation (14).

### Quantifying picornavirus binding to bacterial cells

Bacterial binding assay was performed as previously described for poliovirus (14). Briefly, approximately 3,000 CPM (approximately 1 × 10^6^ PFU) of ^35^S-radiolabeled virus was mixed with PBS+ or 1 × 10^8^ CFU of bacteria and incubated at 37°C in presence of CO_2_ for 1 h. After incubation, bacteria was pelleted and washed with PBS+ to remove unbound virus. The amount of CPM (virus bound to bacterial cells) was determined by scintillation counting.

### Quantifying effects of bacteria on virion stability

To determine the effect of bacteria on thermal stability of picornaviruses, 1 × 10^5^ PFU of each virus was mixed with PBS+, 1 mg/mL of bacterial surface polysaccharides, or 1 × 10^10^ CFU of bacteria and incubated at 44°C for 4.5 h. The same procedure was followed for elevated temperature assays. After incubation, plaque assays were performed using virus-specific conditions to determine the amount of viable virus before and after heat treatment.

Bleach inactivation assay was performed as previously described for PV, except that a lower concentration of bleach was used here (7). Briefly, 1 × 10^5^ PFU of each virus was mixed with PBS+, 1 mg/mL of bacterial surface polysaccharides, or 1 × 10^8^ CFU of bacteria. Samples were incubated at 37°C for 1 h and then added to 0.0001 % fresh bleach for 1 min. Bleach neutralization was done by adding 0.01 % sodium thiosulfate (Sigma). Plaque assays using virus-specific conditions were performed to determine amount of viable virus before and after bleach treatment.

To examine effects of feces on viral stability, feces from four-to ten-week-old male C57BL/6 *PVR-IFNAR -/-* mice were collected and resuspended in PBS+ to a final concentration of 0.0642 mg/mL. Briefly, 1 × 10^5^ PFU of virus was mixed with 300 mL of PBS+ or resuspended fecal samples and incubated at 37°C in the presence of 5 % CO_2_. Additional samples in PBS+ were placed at 4°C as a control. Samples were taken at designated time points and processed by chloroform extraction as previously described (3,7). Plaque assay was performed to determine amount of viable virus before and at designated time points, as described earlier. In Figure 3F, 1 × 10^5^ PFU of Mengo virus was mixed with approximately 1 × 10^5^ CFU each of *E. coli, P. ruminicola*, and *L. johnsonii* in a total volume of 300 mL and samples were incubated at 37°C. Samples were collected and titers were determined as described above.

### Mouse Experiments

Animals were handled according to the Guide for the Care and Use of Laboratory Animals. C57BL/6 *PVR-IFNAR -/-* mice were obtained from S. Koike (Tokyo, Japan) (31). Feces collection was performed at UT Southwestern Medical Center.

### Data Analysis

Figures of viral structures were generated using the UCSF Chimera software (http://www.rbvi.ucsf.edu/chimera). The Electron Microscopy Data Bank (EMDB) IDs used for each virus are as follows: Aichi virus (EM-9517), CVB3 (EM-6637) and Saffold virus (EM-3097) to represent their respective genera. The phylogenetic tree was generated using the MEGA7 software and following the Neighbor-Joining Method (32). The optimal tree with the sum of branch length = 2.65090397 is shown. The tree is drawn to scale, with branch lengths in the same units as those of the evolutionary distances used to infer the phylogenetic tree. The evolutionary distances were computed using the Poisson correction method and are in the units of the number of amino acid substitutions per site. (32). The analysis involved 5 amino acid sequences. All positions containing gaps and missing data were eliminated. There were a total of 766 positions in the final phylogeny tree dataset.

All statistical analyses were performed using the GraphPad Prism Software. Outliers were identified and removed by the ROUT method, Q = 1%. All one way ANOVA tests were performed with Dunnett’s multiple comparisons post hoc test. All two-way ANOVA tests were performed with Tukey’s post hoc test.

## ACKNOWLEDGEMENTS

We thank Andrea Erickson, Broc McCune and Arielle Woznica for critical review of the manuscript. We thank Marco Vignuzzi for the Mengo virus infectious clone. We also thank Mikal Woods-Acevedo for providing ^35^S-labeled CVB3-H3 and CVB3-Nancy viruses. We also thank Nam Nguyen for assistance with structural modeling. Molecular graphics and analyses were performed with the UCSF Chimera package. Chimera is developed by the Resource for Biocomputing, Visualization, and Informatics at the University of California, San Francisco (supported by NIGMS P41-GM103311).

